# Synthetic EMG Based on Adversarial Style Transfer can Effectively Attack Biometric-based Personal Identification Models

**DOI:** 10.1101/2022.10.14.512221

**Authors:** Peiqi Kang, Shuo Jiang, Peter B. Shull

**Author notes:** Corresponding authors: Shuo Jiang and Peter B. Shull.

## Abstract

Biometric-based personal identification models are generally considered to be accurate and secure because biological signals are too complex and person-specific to be fabricated, and EMG signals, in particular, have been used as biological identification tokens due to their high dimension and non-linearity. We investigate the possibility of effectively attacking EMG-based identification models with biological adversarial input via a novel EMG signal individual style transformer based on a generative adversarial network. EMG hand gesture data from eighteen subjects and three well-recognized deep EMG classifiers were used to demonstrate the effectiveness of the proposed attack methods. The proposed methods achieved an average of 99.41% success rate on confusing identification models and an average of 91.51% success rate on manipulating identification models. These results demonstrate that EMG classifiers based on deep neural networks can be vulnerable to synthetic data attacks. The proof-of-concept results reveal that synthetic EMG biological signals must be considered in biological identification system design across a vast array of relevant biometric systems to ensure personal identification security for individuals and institutions.

## I. Introduction

ELECTROMYOGRAPHY (EMG)-based hand gesture recognition is a representative application of human-machine interface technology [1]. Due to the high information related to neural activities and high time resolution, EMG-based hand gesture recognition methods have played vital roles in prosthetic control, VR/AR interaction, user verification, and identification [2].

Since the EMG signals are high dimensional and generated from complex physiological structures, the EMG signals of the same hand gesture have significant differences between individuals. These individual differences will pose technical challenges on applications including prosthetic control but in turn hold potential to be reliable tokens for personal identification. With the help of machine learning or deep learning recognition models, EMG-based user identification systems can achieve a 1% to 4% averaged equal error rate using multi-channel EMG bands or high-dimensional EMG systems [3], [4].

These research studies show that EMG-based user identification systems are highly functional, and since biological signals are high dimensional and complex, biometric-based identification systems are usually considered to be highly secured. However, in recent years, many attack methods have been proposed and received intensive attention; these methods usually add designed small perturbations into the input data to manipulate models’ output or make models confused [5]. A brief literature review on attacking biologic recognition models will also be provided in the next section. Among these models, the most notable is the deployment of generative neural networks in image synthesis [6]. These synthetic images successfully cheated the most state-of-the-art identification models and caused huge security risks. However, for attacking EMG-based identification models using synthetic EMG signals, there are few related studies, but these attacks may cause huge security problems and potentially inevitable losses to users.

Therefore, we proposed two assumptions in this paper: first, an individual’s EMG style can be learned by a neural network; second, with the learned EMG style, synthetic EMG signals with the same style can be generated and used to attack EMG-based identification models. To validate these assumptions, we built an EMG signal style transformer based on the cycle generative adversarial network to learn an individual’s EMG hand gesture signal style and generate other individuals’ synthetic EMG hand gesture signals in the same style (Fig. 1). Experiments on three deep identification models and eight hand gestures of eighteen subjects validated that the artificial synthetic EMG signals can effectively cheat deep identification models and manipulate them to achieve the attacker’s desired outcome. It is worth mentioning that, due to the high dimension and non-linearity of EMG signals, no two EMG segments should be the same; otherwise, they will be easily identified as synthetic attacks by the security system. In addition, when encountering identification systems requiring long-time sampling, few leaked data will soon run out, so attackers will be forced to use repeated data and thus be detected. Therefore, using leaked data to directly attack identification systems is not practical and deep fake data generation methods are necessary. In addition, the focus of this paper lies in exploring the possibilities of potential attacks from the biological perspective instead of from the mathematical perspectives that aim at the working principle of identification models.

**Fig. 1.**
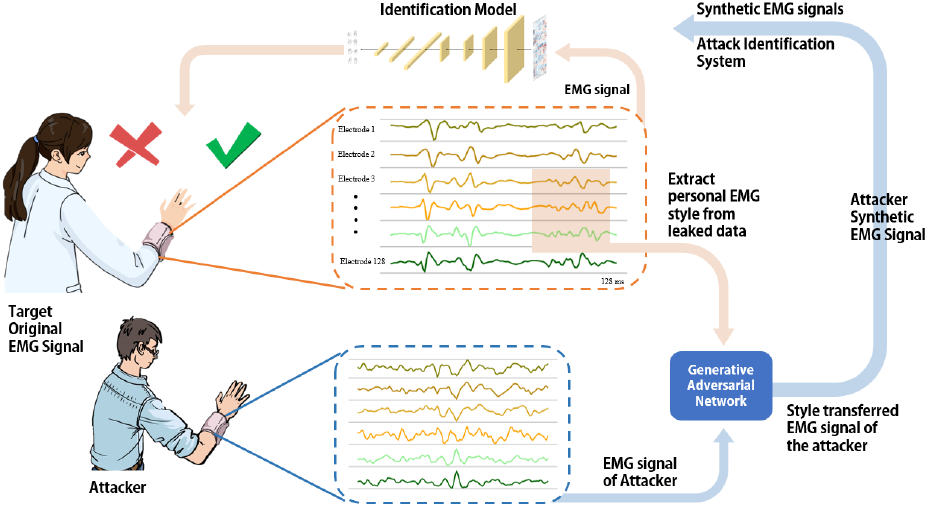
Suppose that an identification model based on a deep convolutional neural network (CNN) uses EMG hand gesture signals input from a 128-channel high-density EMG armband as tokens. However, if a user’s EMG signals were leaked due to various reasons, his personal EMG style can be extracted from them and used as a target style to generate synthetic EMG signals via a generative adversarial network. With these synthetic EMG signals, attackers can effectively hack the identification system and manipulate its outputs, which may cause huge risks to personal privacy and information security.

**Fig. 2.**
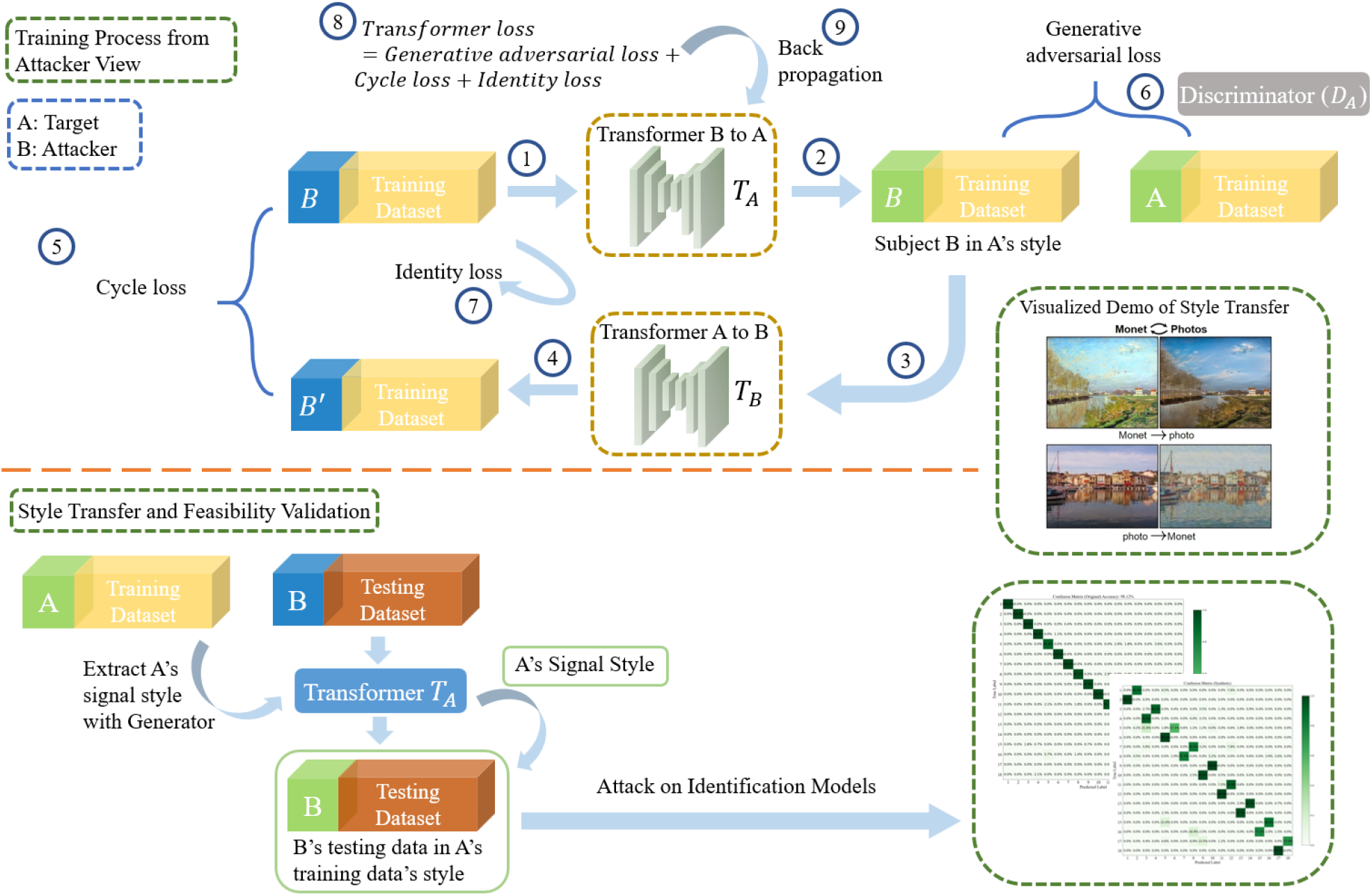
The training process of the proposed signal transformer from B’s (attacker) view. Suppose there are two subjects: A and B. To train a signal transformer that can generate subject B’s EMG signal in subject A’s style, the transformer should be optimized in the gradient descent direction of the sum of the generative adversarial loss, cycle consistency loss, and identity loss. The aim and function of the three kinds of loss are defined and introduced in the Methods section. The figure on the right shows a visualized demo of the style transfer algorithm. After the training is finished, new data from B can be transferred into A’s style and be used in the attack on personal identification models.

The contributions of this paper are as follows: (1) We presented a novel method for generating style-transferred synthetic EMG hand gesture signals. (2) We demonstrated that synthetic EMG signals are capable of cheating the personal identification systems which implies that the synthetic biological signals must be taken into consideration in biological identification systems, design. Our methods hold the potential to be used to guide the design of millions of identification devices to protect people’s privacy and information safety.

This paper was organized as follows: in Section II, a brief literature review on related studies is presented. In Section III, the problem formulation and structure of the proposed synthetic signal generation methods are introduced. In Section IV, experimental protocol, validation protocol, and results are introduced. Finally, in Section V, discussion on our attack methods and results are presented.

## II. Background Research

### A. EMG-based User Identification

Recently, various novel identification technologies based on biometric technologies have been reported by worldwide researchers. Since the gesture-recognition-based identification technologies using EMG signals can be collected during human hand activity and are similar to the process of traditional identification methods (e.g., entering passwords by hand), they have been intensively explored by various researchers. Yamaba et al. [7] proposed the idea of utilizing EMG signals for user identification and achieved promising results using a support vector machine (SVM) classifier. Jiang et al. [8] utilized HD-sEMG signals of common daily hand gestures as identification inputs and proposed a cancelable HD-sEMG-based biometrics system to protect personal information security. Pradhan et al. [9] designed a series of experiments on the effect of different feature extraction methods and the number of channels to the EMG-based identification system and systematically investigated the performance of sixteen static wrist and hand gestures.

Compared with other biometric identification systems (e.g., gait-based [10], electrocardiograph (ECG)-based [11], and electroencephalograph (EEG)-based [12]), the EMG-based methods show advantages on high information security, high signal-to-noise ratio, high recognition accuracy, and more convenient acquisition [3].

### B. Attacks on Biological Classifiers

Artificial intelligence models represented by deep learning models achieved remarkable success in various recognition tasks, but their vulnerability to interference or attacks also drew great attention [13]. There are three kinds of attack methods according to how deep the attacker can get access to the target models: white-box attacks, gray-box attacks, and black-box attacks. The black-box attacks are most practical because they only need to know the input and output of the target models. Su et al. [14] successfully cheated image classifiers by changing a single pixel of the input. Recently, the famous Open AI lab announced a simple but highly effective method called the typographic attack: simply pasting a tag with a note on the object can mislead the state-of-the-art recognition models [15]. With a similar strategy, researchers broke the state-of-the-art Face ID system with a printed sticker [16]. Attacks on the brain-computer interface (BCI) were also investigated. Zhang et al. [17] achieved effective attacks on EEG-based BCIs and proved the vulnerability of convolutional neural network (CNN) classifiers under small deliberate perturbations. Liu et al. [18] proposed a total loss minimization approach to generate universal adversarial perturbations to attack EEG-based BCIs and successfully manipulated the output of the models. Zhang et al. [19] and Bian et al. [20] conducted intensive studies on the attack on EEG-based BCI spellers, and the results showed that the BCI spellers can be easily manipulated and may cause serious problems like medical misdiagnose.

### C. Synthetic Biological Signals

The generative neural network is a kind of artificial intelligence model that can generate synthetic images or data, and the generative adversarial network (GAN) is the most representative [21]. GAN is a deep learning frame that consists of a generator and a discriminator. The generator is used to generate synthetic data, and the discriminator is used to discriminate between the synthetic data and real data, therefore helping the generator to generate more realistic synthetic data.

In addition, GAN also provides a new way to generate synthetic biological signals which were considered too complex to generate synthetically. Jiao et al. [22] utilized Wasserstein GAN generated EEG and EOG signals to expand the dataset and improve classifier performance in driver sleepiness detection. Other research also reports the contribution of Wasserstein GAN in emotion recognition [23]. Access to personal ECGs is restricted because of privacy concerns, but building automated computer-aided diagnosis systems requires vast amounts of data. Nankani et al. [24] proposed an approach for generating irregular beats (e.g., supraventricular ectopic, ventricular ectopic, and normal beats) with a conditional GAN to generate synthetic ECG data for diagnosis systems’ datasets. Ding et al. [25] proposed a log-spectral matching GAN to generate PPG signals for atrial fibrillation detection.

Generated synthetic EMG data also received attention from the neuroscience community. Anicet et al. [26] utilized deep convolutional generative adversarial networks and style transfer to generate Parkinson’s disease EMG signals to reduce the displeasure and pain of patients to collect lots of data. Campbell et al. [27] validated the feasibility of generating EMG signals with hand motions information using a deep generative model called sinGAN. Bird et al. [28] utilized a generative model called the generative pre-trained transformer to generate synthetic EMG signals of three gestures, including hand open, hand closed, and at rest, and demonstrated the synthetic data in a prosthetic hand control application.

To our best knowledge, there were no studies focused on EMG style transferring between individuals using generative methods or studies that tried to disclose the security risk of synthetic data in EMG-based identification systems. This work indicated that the resistance to synthetic biological signals and data protection or encryption must be taken into consideration in biological identification systems design. We hope our work can draw researchers’ attention to the potential threat of the synthetic biological signals to personal information security and property security.

## III. Methods

### A. Problem Formulation and Overview

Suppose a biometric-based identification model can accurately recognize a type of EMG hand gesture signals as subject A:

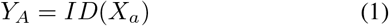

where *ID* is the identification model, *X*_*a*_ is the data input from subject A, and *Y*_*A*_ is the identification result of the identification model. Due to the individual differences in EMG signals between two individuals, if the same model received signals from other subjects, for instance, attacker B, then the system will recognize this subject is not A.

However, if subject A’s EMG data was leaked due to various reasons, this data might be used to attack the identification model. With this leaked data, we proposed a generative adversarial EMG signal transformer to generate synthetic EMG signals to manipulate the identification model’s outputs:

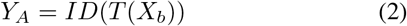

where *T* is the generative adversarial EMG signal transformer we proposed. and *X*_*b*_ is the data input from attacker B. In a real-life scenario, if the transformer model was embedded into an identification system, by transferring the input data into synthetic data, the attacker could manipulate the identification system’s outputs in real-time.

Our proposed signal transformer’s network architecture from top to bottom is a reflection padding layer, three convolutional layers (kernel sizes are seven, three, three), four residual blocks, two convolutional layers (kernel size is three) with an up-sample function (scale factor is two), a reflection padding layer, and a convolutional layer (kernel size is seven) (Fig. 3). The input is 128 ms of EMG signal of 128 channels, and the output size is the same.

**Fig. 3.**
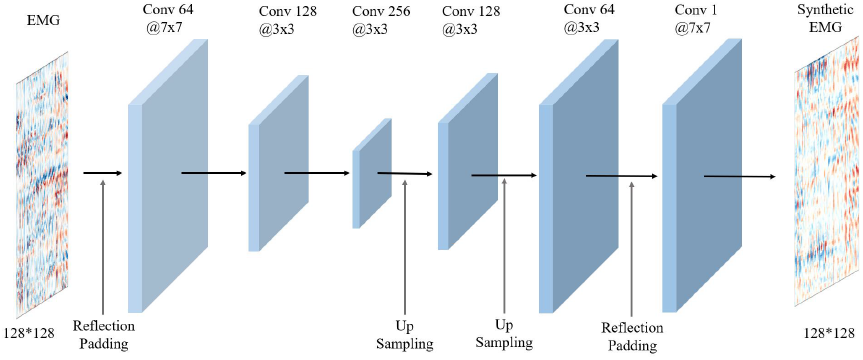
The network architecture of the transformer. The input is a 128 ms of EMG signal of 128 channels, and the output is the same size synthetic EMG signal.

### B. EMG Signal Transformer Training Process

Training the transformer requires leaked data from subject A and data from attacker subject B. During the training process, three kinds of loss were used to guide the optimization direction of the parameters of the transformer.

First, to improve the quality of the synthetic signal, we utilized the idea of generative adversarial networks and designed a discriminator for the training of the signal transformer [29]. The network architecture of the discriminator consists of four convolutional layers (kernel size is four), a zero-padding layer, and a convolutional layer (kernel size is four). During the training process, the synthetic signals generated by the transformer will be scored by the discriminator. The discriminator will give higher scores to synthetic signals that are similar to the leaked data from subject A and give lower scores to synthetic signals that are less similar to the leaked data from subject A. During the training process, the transformer and discriminator will both be trained, and therefore, can continually force the transformer to generate high-quality synthetic signals. The loss used in this process was defined as the generative adversarial loss:

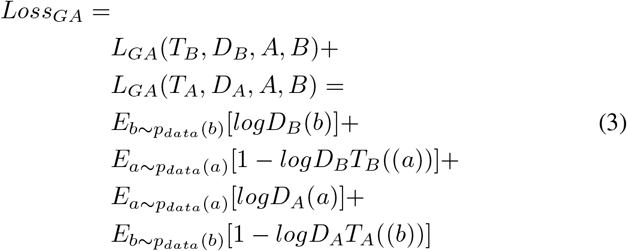

where *A* is the subject whose data was leaked, *B* is the attacker, *a* and *b* are data samples from *A* or *B*, *T*_*A*_ is the transformer that transform *B*’s data into *A*’s style, *T*_*B*_ is the transformer that transform *A*’s data into *B*’s style, *D*_*A*_ and *D*_*B*_ are discriminators that calculate the difference between real data and synthetic data, E is the expected value, and *a* ~ *p*_*data*_(*a*) and *b* ~ *p*_*data*_(*b*) represent data distribution.

Second, in order to transform the data style and keep the content unchanged, we adapted ideas from sentence translation and cycle generative adversarial networks to add an additional constraint condition into the training process of the signal transformer [30]. In sentence translation, the goal is keep the original meaning but translate the sentence into a different language. To make sure the translated meaning is correct, translators always adopt a cycle strategy that translates the translated sentence back to the original language and compares its meaning with the original untranslated sentence and try to make the difference as small as possible. We defined this type of loss as cycle loss. The aim of the *Loss*_*cycle*_ is to force the transformer to generate synthetic data only different from the real data in the style aspect and was defined by:

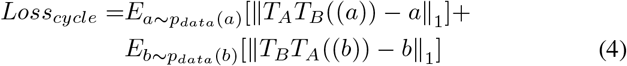

Third, due to the high dimensionality and nonlinearity of EMG signals, no two EMG segments should be the same; otherwise, they will easily be identified as synthetic attacks by the security system. To prevent the transformer from just cheating the discriminator by directly outputting subject A’s leaked data as the transformed B’s synthetic signal, in which case the transformation process would be meaningless, we utilized an identity loss as an additional constraint. The working principle of the *Loss*_*identity*_ is that, if a transformer can generate data in *A* style and if input data is already in *A* style, then the synthetic data should stay the same:

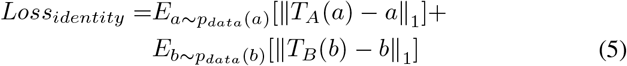

Therefore, the loss of the training process of the generative adversarial EMG signal transformer was given by:

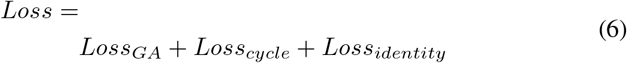

Moreover, since the training of transformer and discriminator is adversarial, when training the transformer, parameters of *D*_*A*_ and *D*_*B*_ should be frozen, and when training the discriminator, parameters of *T*_*A*_, *T*_*B*_ and one of the two discriminators should be frozen. Therefore, the optimization objective of the transformer was:

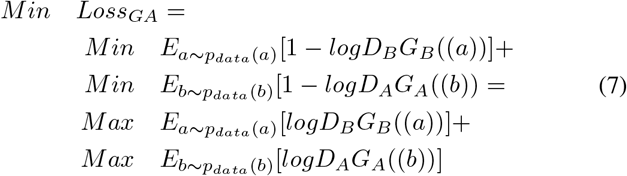

This optimization objective means optimizing parameters of *T*_*A*_ and *T*_*B*_ to get higher scores from *D*_*A*_ and *D*_*B*_ and to force *T*_*A*_ and *T*_*B*_ to generate more realistic synthetic data.

In addition, the discriminator should try to prioritize real data (maximize *D*_*A*_(*a*)) and try to minimize synthetic data (minimize *D*_*A*_(*G*_*a*_(*b*))). Therefore, the optimization objective of one discriminator (e.g., *D*_*A*_) was:

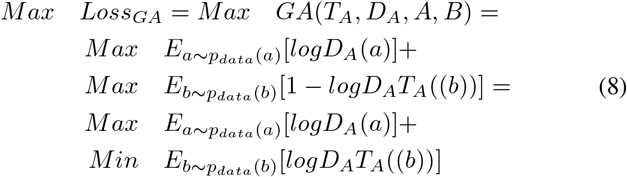

After the transformer was trained, new data from attacker B can be transformed into subject A’s style, and in the next section, we will validate the feasibility of attacking the identification system with the synthetic subject A’s data.

## IV. Experiments and Results

### A. Dataset and Experimental Protocol

CapgMyo dataset A was chosen to validate our proposed methods [31]. The CapgMyo A is a public dataset consisting of eight hand gestures’ HD-EMG records (8×16 electrode array) of 18 participants. Eight hand gestures were included in the dataset: 1. Thumb up (TU); 2. Extension of index and middle, flexion of the others (EIM); 3. Flexion of the ring and little finger, the extension of the others (FRL); 4. Thumb opposing base of the little finger (TO); 5. Abduction of all fingers (AA); 6. Fingers flexed together in the first (FF); 7. Pointing index (PI); 8. Adduction of extended fingers (AE). The signals were sampled at 1,000 Hz, filtered with a band-pass filter at 20-380Hz, and normalized to the [−1, 1] range, corresponding to the voltage of [−2.5 mV, 2.5 mV]. Each gesture of each subject had ten trials. To ensure the reliability of experiment results, the odd-numbered trials of each subject were used as the training dataset and the even-numbered trials of each subject were used as the testing dataset.

### B. Identification Models and Baseline Benchmark

It is vital to select powerful identification models as attack targets; otherwise, the results can’t prove the effectiveness of the attack methods because good attack results might be caused by weak baseline models. Therefore, for the reliability of our proposed methods, we selected three well-recognized deep EMG classification models to use as identification models and target with attacks: GengNet [31], EMGNet [32], and VGG16Net [33]. We trained these identification models with eighteen subjects’ training datasets and tested them with eighteen subjects’ testing datasets. For the performance evaluation, we selected the rank-k identification rate as the evaluation metric and calculated the cumulative match characteristic (CMC) curve. In addition, as the commonly used metrics, the values of rank-1 and rank-5 were also listed, and the confusion matrixes were provided. These evaluation metrics are representative and well-recognized quantification methods for evaluating biometric-based identification models. To make sure that our result is robust to different hand gestures, for each gesture, we trained three different structured identification models. If not specifically mentioned, all the results present in the following are the mean values of all eight gestures. The test results of three identification models were shown as follows: the rank-1 and rank-5 results showed that our models have strong identification ability (Table. I), the CMC curve showed that identification systems based on these models will have reliable identification ability in a real-life scenario (Fig. 6), and the confusion matrixes showed that our identification models have good robustness among different subjects (Fig. 5). These results demonstrated that our identification models are powerful and qualified to be attack targets.

**Fig. 4.**
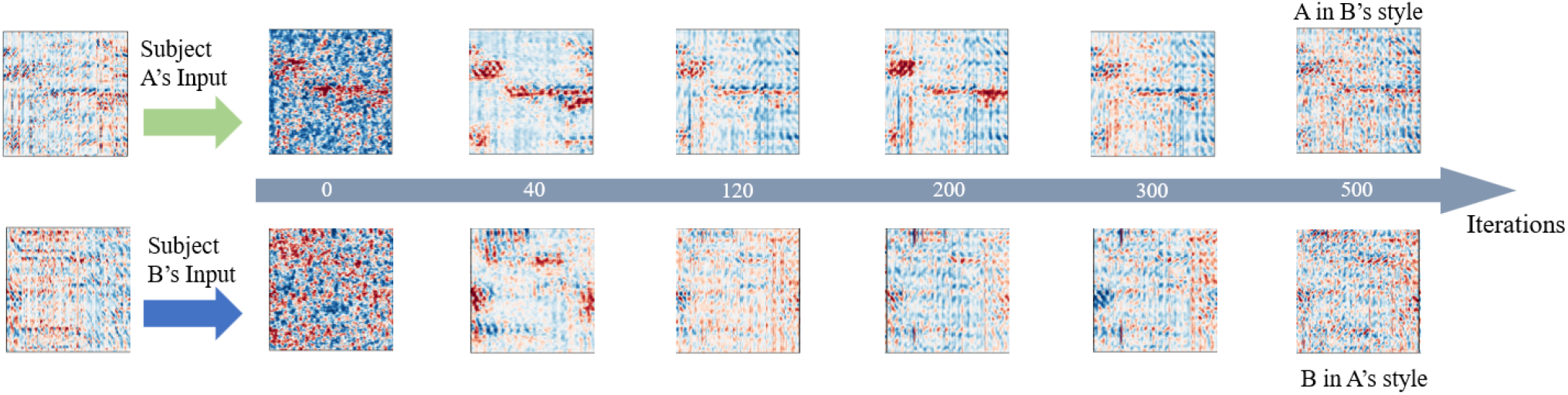
Evolution of the synthetic signals during the training process. When the training started, the synthetic signals were basically irregular noises. As the training proceeded, the synthetic signals learned features from the original signals but still had major defects in some areas. After a certain number of iterations, the synthetic signals became indistinguishable from the original signals, and the value of the loss function almost stopped decreasing at which point the training could be considered finished.

**Fig. 5.**
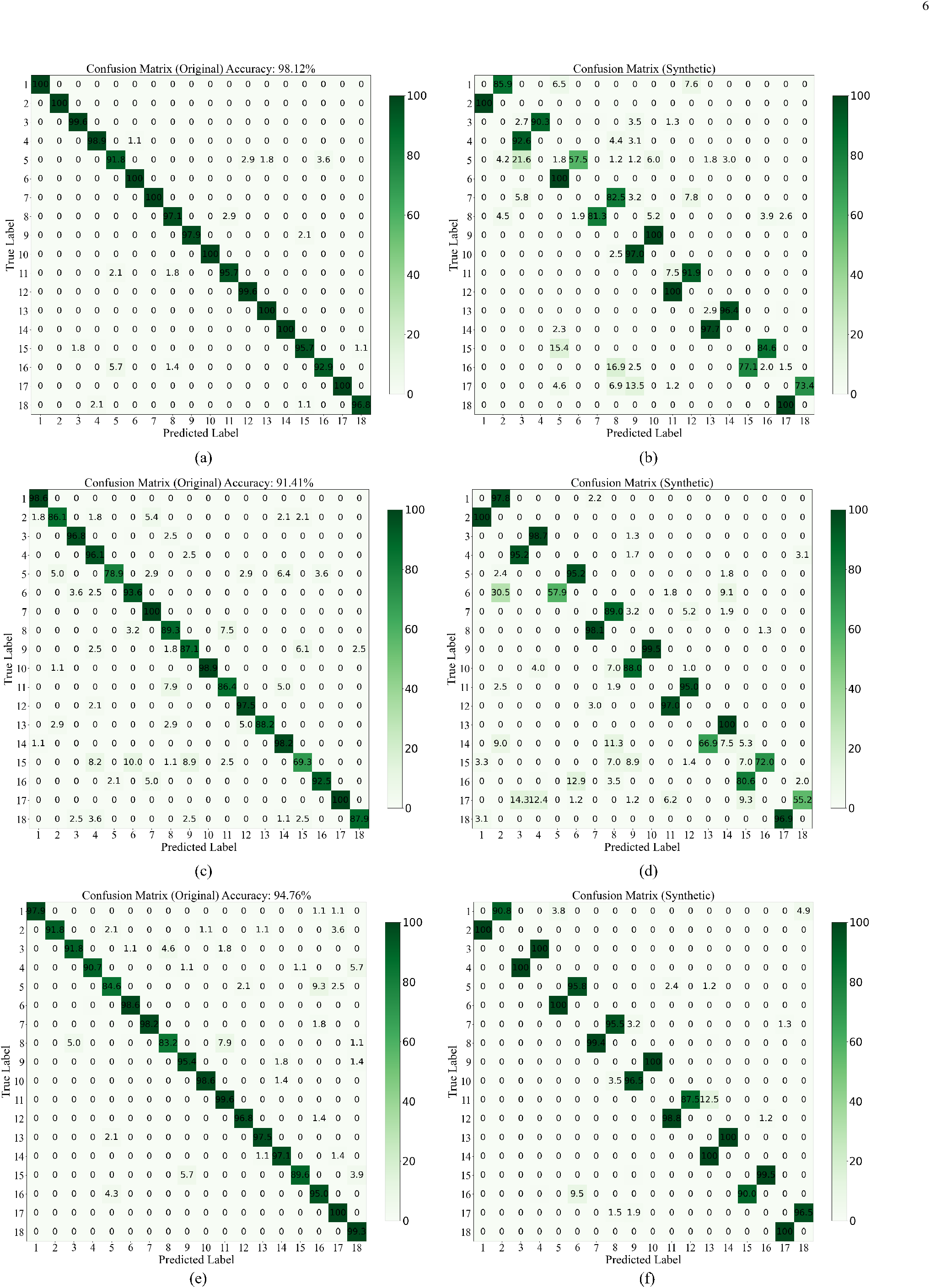
The data are averaged across eight hand gestures. (a) Confusion matrix for eighteen subjects’ identification when testing on original data using the GengNet. (b) Confusion matrix for eighteen subjects’ identification when testing on synthetic data using the GengNet. (c) Confusion matrix for eighteen subjects’ identification when testing on original data using the EMGNet. (d) Confusion matrix for eighteen subjects’ identification when testing on synthetic data using the EMGNet. (e) Confusion matrix for eighteen subjects’ identification when testing on original data using the VGG16Net. (f) Confusion matrix for eighteen subjects’ identification when testing on synthetic data using the VGG16Net. The (a), (c), and (e) confusion matrices aim to validate the strong identification ability of the GengNet, EMGNet, and VGG16Net. The (b), (d), and (f) confusion matrices aim to demonstrate the effectiveness of attacking GengNet, EMGNet, and VGG16Net with synthetic data.

**Fig. 6.**
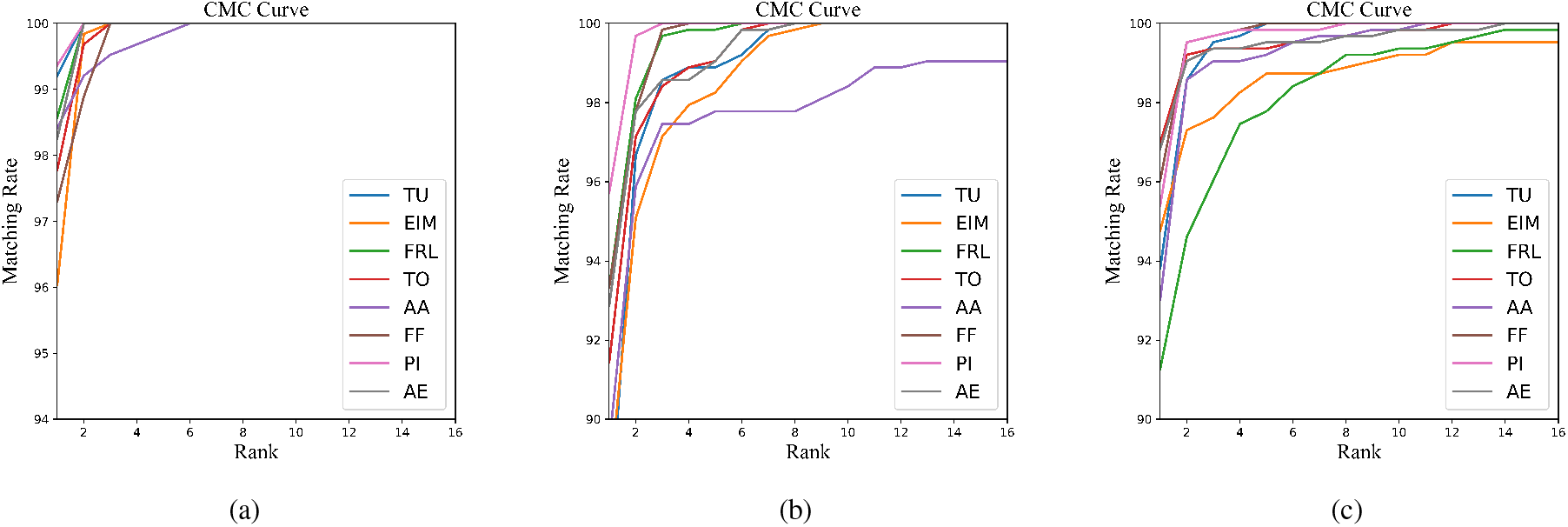
The data are averaged across eighteen subjects. These figures showed that our identification models are powerful and qualified to be attack targets. (a) CMC curves for each gesture of the GengNet-based identification model. (b) CMC curves for each gesture of the EMGNet-based identification model. (c) CMC curves for each gesture of the VGG16Net-based identification model.

**Table 1.**
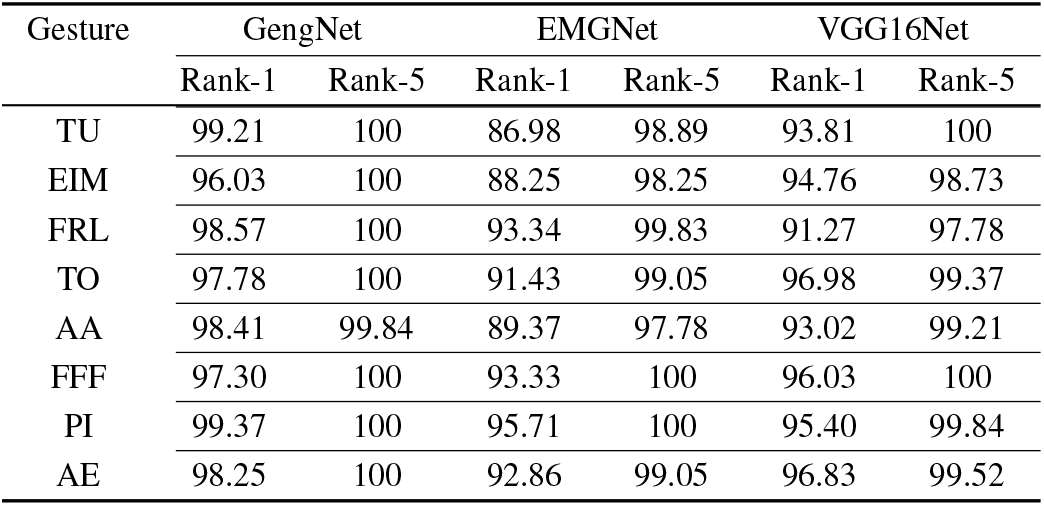
RANK-1 AND RANK-5

### C. Synthetic Signal Generation and Attack on Identification Models

Two adjacent individuals were paired with one acting as the data leaker and the other as the attacker to validate the proposed EMG signal transform attack methods. Therefore, nine pairs (eighteen individuals) were tested for each experimental condition. For better understanding, odd-numbered subjects were named as subject A and even-numbered subjects were named as subject B. During the training process, 90% of odd-numbered trials’ data of subjects A and B were used as training data for the transformers, and the other 10% of odd-numbered trials’ data were used as validation data to calculate the loss of the transformers. The training process repeated until the validation loss met the requirement. After the training was finished, we used subject A’s and subject B’s even-numbered trials’ data separately to generate subject A’s EMG signals in B’s style and subject B’s EMG signals in A’s style. We also visualized the signal generation result of transformers at different training stages as shown in Fig. 4.

After the synthetic signals were generated, we tested the identification models with synthetic EMG signals (style transferred between neighboring subjects) to simulate potential attacks. Results showed that the artificial synthetic EMG signals can efficiently cheat the three kinds of identification models in all eight gestures and mislead them to make judgments as we designed. To quantifiably evaluate the proposed attack methods, we defined two evaluation metrics called hit rate and confusion rate, which represent the success rate of using synthetic EMG signals on attacking identification models and the chance of identification models being confused by the synthetic EMG signals:

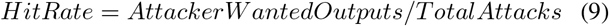

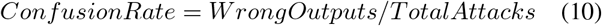

Wrong outputs included outputs that were what the attacker wanted and outputs that were not what the attacker wanted but were still not correct.

Under attacks of the synthetic EMG signals, identification models’ rank-1 and rank-5 evaluation metrics dropped close to zero, which means that these identification models were disabled. The hit rate of the attack methods on three different identification models (GengNet, EMGNet, and VGG16Net) were 89.34%, 87.94%, and 97.24%, respectively. The confusion rate of the attack methods on three different identification models (GengNet, EMGNet, and VGG16Net) were 99.06%, 99.17%, and 100%, respectively. These results showed that our attack methods can effectively manipulate and confuse the state-of-the-art identification models (Table II).

**Table 2.** HIT RATE AND CONFUSION RATE

| Gesture | GengNet |  | EMGNet |  | VGG16Net |  |
| --- | --- | --- | --- | --- | --- | --- |
|  | Hit | Confusion | Hit | Confusion | Hit | Confusion |
| TU | 99.20 | 99.61 | 94.16 | 100 | 97.27 | 100 |
| EIM | 93.82 | 100 | 89.58 | 99.31 | 100 | 100 |
| FRL | 83.84 | 100 | 88.04 | 96.43 | 96.61 | 100 |
| TO | 98.29 | 100 | 82.66 | 100 | 98.81 | 100 |
| AA | 76.62 | 97.22 | 88.43 | 97.42 | 99.37 | 100 |
| FFF | 80.88 | 97.23 | 90.89 | 100 | 95.67 | 100 |
| PI | 95.66 | 100 | 84.74 | 98.77 | 95.64 | 100 |
| AE | 98.84 | 100 | 90.68 | 100 | 99.38 | 100 |

## V. Discussion

In this paper, we proposed a novel personal identification model attack methods based on the EMG signal transformer. Experiments on eighteen subjects and three strong identification models proved its feasibility and reliability. Our methods achieved an average of 99.41% success rate on confusing identification models and an average of 91.51% success rate on manipulating identification models. These results demonstrate that our methods hold the potential to be used to guide the design of millions of identification devices to protect people’s privacy and information safety.

Using the same training and testing conditions, of all the models the GengNet had the strongest identification ability. Compared with the other two identification models, the GengNet has two conventional layers whose kernel size is one and much fewer channel numbers. This result shows that with a smaller kernel size identification models can obtain stronger recognition ability with fewer channels. However, smaller kernel size models will require a significantly longer time and more computing resources to train. Therefore, if the accuracy has already met the requirement, a lightweight model might also be an option.

In addition, results also showed that the anti-attack ability has no direct relationship with the identification ability. Three models with different structures show no significant difference (Tukey’s test based on analysis of variance, *p* < 0.05) in confusion rate, which indicated that our attack methods have strong universality. We think the performance difference shown in the hit rate might be caused by the different generalization abilities of models. Models with high generalization ability tend to have a lower hit rate and models with low generalization ability got a higher hit rate. Therefore, we think improving the generalization ability of identification models might improve the ability to resist target attacks but still can’t avoid being confused. These results indicate that the best way to improve identification systems’ anti-attack ability, apart from developing a more robust identification model, we might need to take the system of view. Applying more complex encoding strategies or adopting sensor fusion technologies may also get a better result. What is more important is to protect original data from leaking, and some studies have already proposed using encrypted data to improve data security.

For style transferred signal generating, generative models are height unconstrained and very difficult to train. During the training process, the loss will randomly uprush, and the time consumption is also larger than training a normal neural network. In addition, the signal transformer has multiple solutions, and any transformers that meet loss requirements can be used. Therefore, setting an appropriate loss threshold will accelerate the training process. However, there is no clear standard on the loss value. A recommended method is to visualize synthetic data during the training process and choose an appropriate loss threshold with multiple experiments.

Our proposed EMG signal transferring and generating methods can also be applied in other areas. Since our methods can significantly reduce the individual difference between subjects, it can also be applied in hand gesture recognition or prosthetic control to improve the robustness and recognition accuracy. It can also provide a new solution to improve the capability of the human-computer interface apart from self-adaptation algorithms, transfer learning algorithms, and sensor fusion algorithms.

## VI. Conclusion

This paper is the first work that focused on EMG hand gesture style transferring between individuals and the potential risk of synthetic data in personal identification systems. Rigorous experiments proved the feasibility of the cycle generative adversarial network based EMG style transformer and the vulnerability of deep EMG classifiers. The result demonstrated that synthetic biological signals must be taken into consideration in any biological identification system design, and the protection of biological signals, including EMG signals, is vital for personal privacy and security.

## Notes

This work was supported by the National Natural Science Foundation of China under Grant 51875347 and Grant 52105033, Shanghai Municipal Science and Technology Major Project under Grant 2021SHZDZX0100, and Chenguang Program by Shanghai Municipal Education Commission under Grant 21CGA23.

### Competing Interest Statement

The authors have declared no competing interest.

